# Village in a dish: a model system for population-scale hiPSC studies

**DOI:** 10.1101/2021.08.19.457030

**Authors:** Drew R. Neavin, Angela M. Steinmann, Han Sheng Chiu, Maciej S. Daniszewski, Cátia Moutinho, Chia-Ling Chan, Mubarika Tyebally, Vikkitharan Gnanasambandapillai, Chuan E. Lam, Uyen Nguyen, Damián Hernández, Grace E. Lidgerwood, Alex W. Hewitt, Alice Pébay, Nathan J. Palpant, Joseph E. Powell

## Abstract

The mechanisms by which DNA alleles contribute to disease risk, drug response, and other human phenotypes are highly context-specific, varying across cell types and under different conditions. Human induced pluripotent stem cells (hiPSCs) are uniquely suited to study these context-dependent effects, but to do so requires cell lines from hundreds or potentially thousands of individuals. Village cultures, where multiple hiPSC lines are cultured and differentiated together in a single dish, provide an elegant solution for scaling hiPSC experiments to the necessary sample sizes required for population-scale studies. Here, we show the utility of village models, demonstrating how cells can be assigned back to a donor line using single cell sequencing, and addressing whether line-specific signaling alters the transcriptional profiles of companion lines in a village culture. We generated single cell RNA sequence data from hiPSC lines cultured independently (uni-culture) and in villages at three independent sites. We show that the transcriptional profiles of hiPSC lines are highly consistent between uni- and village cultures for both fresh (0.46 < R < 0.88) and cryopreserved samples (0.46 < R < 0.62). Using a mixed linear model framework, we estimate that the proportion of transcriptional variation across cells is predominantly due to donor effects, with minimal evidence of variation due to culturing in a village system. We demonstrate that the genetic, epigenetic or hiPSC line-specific effects on gene expression are consistent whether the lines are uni- or village-cultured (0.82 < R < 0.94). Finally, we identify the consistency in the landscape of cell states between uni- and village-culture systems. Collectively, we demonstrate that village methods can be effectively used to detect hiPSC line-specific effects including sensitive dynamics of cell states.

## Introduction

Using human induced pluripotent stem cells (hiPSCs) and their derivatives to study human complex traits such as diseases and drug response is becoming a new research frontier through the intersection with population genetic approaches^1–4^. hiPSCs are karyotypically normal, self-renewable cells that are generated by reprogramming human somatic cells. They have the ability to differentiate into virtually any cell type in the human body^5^, providing a model system to study human cell types *in vitro*. Recent work has demonstrated that hiPSCs are a powerful system for investigating large-scale inter-individual variation and context-dependent effects that would be challenging to recreate *in vivo*. Here, we consider context-dependent effects to be genetic relationships with phenotypes that are only detectable under specific conditions. For example, some expression quantitative trait loci (eQTLs) are only detected in specific tissues^6^, cell types^2,3,7^, cell states^8^, or following drug^9^ or chemokine^10^ exposure. While hiPSCs are a powerful model system to interrogate these context-specific effects, large-scale hiPSC culture is expensive and time-consuming, creating challenges for studies that require hundreds to thousands of donor lines.

To mitigate some of these limitations, recent studies have applied a village approach to culture hiPSCs - where multiple unrelated lines are cultured and differentiated in a single dish^2,8,11^. One of these studies paired flow sorting of survival motor neuron (SMN) protein levels with whole genome sequencing. The proportion of each hiPSC line in each SMN flow-sorted group of cells were then estimated with computational methods. The study design provided statistical power to detect SMN protein quantitative trait loci (pQTLs) - genetic variants that are associated with protein expression levels of cell lines. While this approach is effective, it is not easily scalable for high-throughput assays of molecular phenotypes. Other studies have applied village culture methods and assayed them with single cell RNA-sequencing (scRNA-seq). In these scenarios, each cell is assigned to a hiPSC line in the pool using demultiplexing methods^12–15^. However, the impact on molecular phenotypes such as gene expression due to the presence of multiple cell lines in a single village culture has not been assessed.

This is a challenge, as it is not clear whether cell signaling in villages will influence the transcriptional profiles of each independent hiPSC line. If such effects were to exist, they would likely lead to biases in the identification of both eQTLs and context-dependent effects due to the creation of ‘artificial’ correlations in the phenotypes between donor lines. Here, we develop village culture systems, and demonstrate their efficacy for population-scale stem cell studies. In particular, we investigate how growth rates impact the proportions of cells from each line in the village, and whether cell signaling alters the transcriptional profiles of individual cells in village culture conditions. We similarly evaluated these properties across multiple independent laboratories.

We show that the inter-line gene expression is unaffected by culturing hiPSC lines in a village or between sites. In other words, the genetic, epigenetic or line-dependent effects that underlie variation in gene expression between different lines is consistent when the cells are cultured separately or in a village. We demonstrate that hiPSC line-specific effects that change across a cell-state pseudotime (dynamic effects) can be reproducibly detected, further supporting the use of village culture systems for large scale studies with hiPSCs. Collectively, our results demonstrate that the village model can be effectively used to detect eQTLs and other cell line specific effects and we provide important details that will allow these approaches to be easily implemented in other laboratories.

## Results

### Experimental and analytical framework

We implemented a multi-phased experimental design that enabled the interrogation of village culture conditions while also comparing inter-laboratory and cryopreservation effects using scRNA-seq (**Figure 1a**).

**Figure 1:**
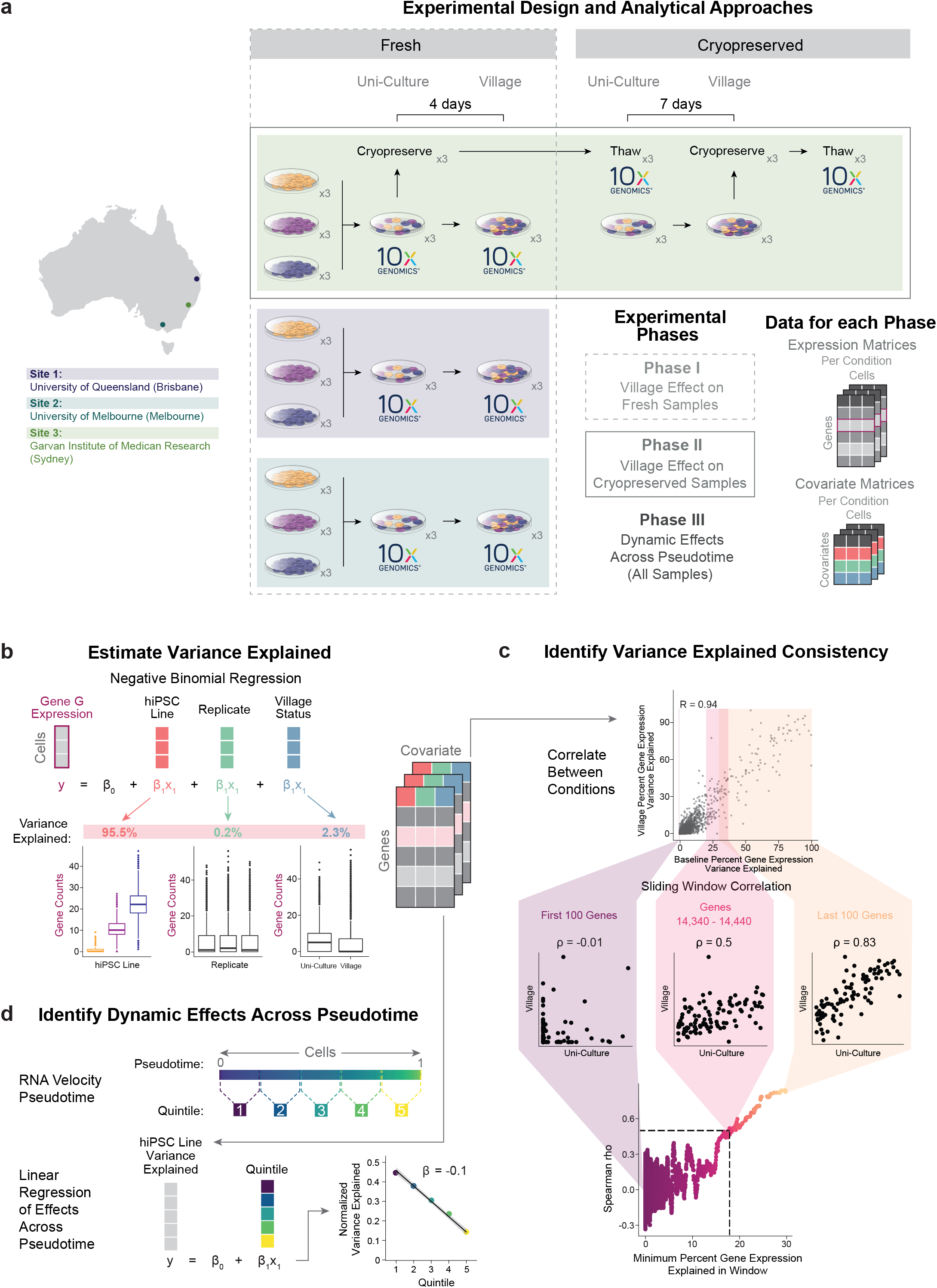
Experimental Design and Analytical Approachesd. **a)** The experimental design to test for the impact of village culture conditions on individual hiPSC lines using scRNA-seq. Phase I compares the impact of the village culture system using fresh samples collected at each site. Phase II investigates whether cryopreservation of village samples impacts individual hiPSC lines. Phase III utilizes all samples to investigate dynamic effects of the hiPSC lines across pseudotime. Each phase utilizes expression matrices that are separated by condition (site for phase I, cryopreservation status for phase II and pseudotime for phase III) as well as covariate matrices for each condition that contain the hiPSC line, replicate and village status. **b)** A negative binomial model is used to estimate the variance of expression for each gene that is explained by each covariate. Those estimates are calculated for each condition in each Phase of the experimental design and used for downstream analyses. **c)** Correlation is used to assess the consistency of the variance explained between different conditions. Then, to further interrogate the point at which the relationship between the two conditions is more consistent, we correlate the variance explained in windows of 100 genes and slide the window by one across all of the genes until the last 100 genes. Those correlation coefficients are then plotted to demonstrate trends in the correlations and identify the points at which a consistent increase in the correlations are achieved. **d)** The pseudotime is estimated for the cells from all samples and they are then partitioned into quintiles. Then, the dynamic effects of the variance explained by the hiPSC lines across the quintiles is detected with a linear model. x3: samples in triplicate; 10x Genomics: 10x scRNA-seq capture.

To compare the effects of village culture conditions in different laboratories, Phase I involved the generation of data from three independent sites - the University of Queensland (Brisbane, Australia; site 1), the University of Melbourne (Melbourne, Australia; site 2) and the Garvan Institute of Medical Research (Sydney, Australia; site 3). The same hiPSC lines from the same passage (FSA0006, MBE1006 and TOB0421)^20^, the same protocols and the same reagents (from the same batches) were used at each site with one exception - hiPSC lines were plated at a lower density at site 3 (∼1/10 the plating density used at sites 1 and 2). Villages were generated using equal proportions of each uni-cultured hiPSC line (cultured in separate dishes) and maintained for four days prior to captures. For the scRNA-seq capture of uni- and village cultures in Phase I, cells were detached and dissociated at the same time at each site and placed on ice. Samples from sites 1 and 2 were transported to site 3 where the samples were processed and captured together (**Figure 1a**; see **Methods** for additional details) - thereby mitigating capture batch effects. Phase II investigated the potential impact of cryopreservation on inter-line effects in the village culture system (maintained for seven days) which was performed at site 3 alone. (**Figure 1a**). Finally, phase III used data from all cells to investigate dynamic hiPSC line effects across cell-state pseudotime (inter-line effects that change over pseudotime; **Figure 1a**).

In the analysis of the scRNA-seq data, we estimate the variance explained by hiPSC line, replicate, and village status using a negative binomial model (**Figure 1b**). The consistency of the variance explained by the hiPSC lines (*i*.*e*., inter-line effects) is then tested by correlating the estimates from uni-culture and village samples (**Figure 1c**). To determine the point at which a high degree of consistency is achieved between samples, we used a sliding window correlation to assess the relationship between every window of 100 genes starting with the 100 genes with the lowest inter-line variation and sliding the window by one gene until the 100 genes with the highest inter-line variation were correlated. Finally, for phase III, the variance of gene expression explained by the hiPSC lines is then tested for dynamic effects across pseudotime with a linear model (*i*.*e*., inter-line effects that change over pseudotime; **Figure 1d**).

### Impact of village culture system

To investigate the potential impacts of village culture conditions on individual hiPSC lines, we collected samples of the hiPSC lines cultured separately and after four days cultured in a village at the three independent sites as previously described (**Figure 1a** and **2a**; see **Methods** for additional details).

### Proportions of hiPSC lines following village culturing

Village culture systems provide advantages over uni-culture systems, provisional that the majority of the hiPSC lines can be maintained in culture. Therefore, we first compared the proportion of each hiPSC line when they were first combined (uni-culture) to the proportions following culture in a village. After demultiplexing the samples (see **Methods**), we found that all hiPSC lines were present in all samples albeit at different proportions in the village than the uni-culture samples at sites 1 and 2 but not 3. At sites 1 and 2, the village samples had a larger proportion of FSA0006 and a smaller proportion of MBE1006 and TOB0421 (**Figure 2b** and **Table S1**). The different proportions of each hiPSC line at sites 1 and 2 in the village compared to the uni-culture samples suggests that growth rate is an important factor to consider when designing village experiments for long term cell culture.

**Figure 2:**
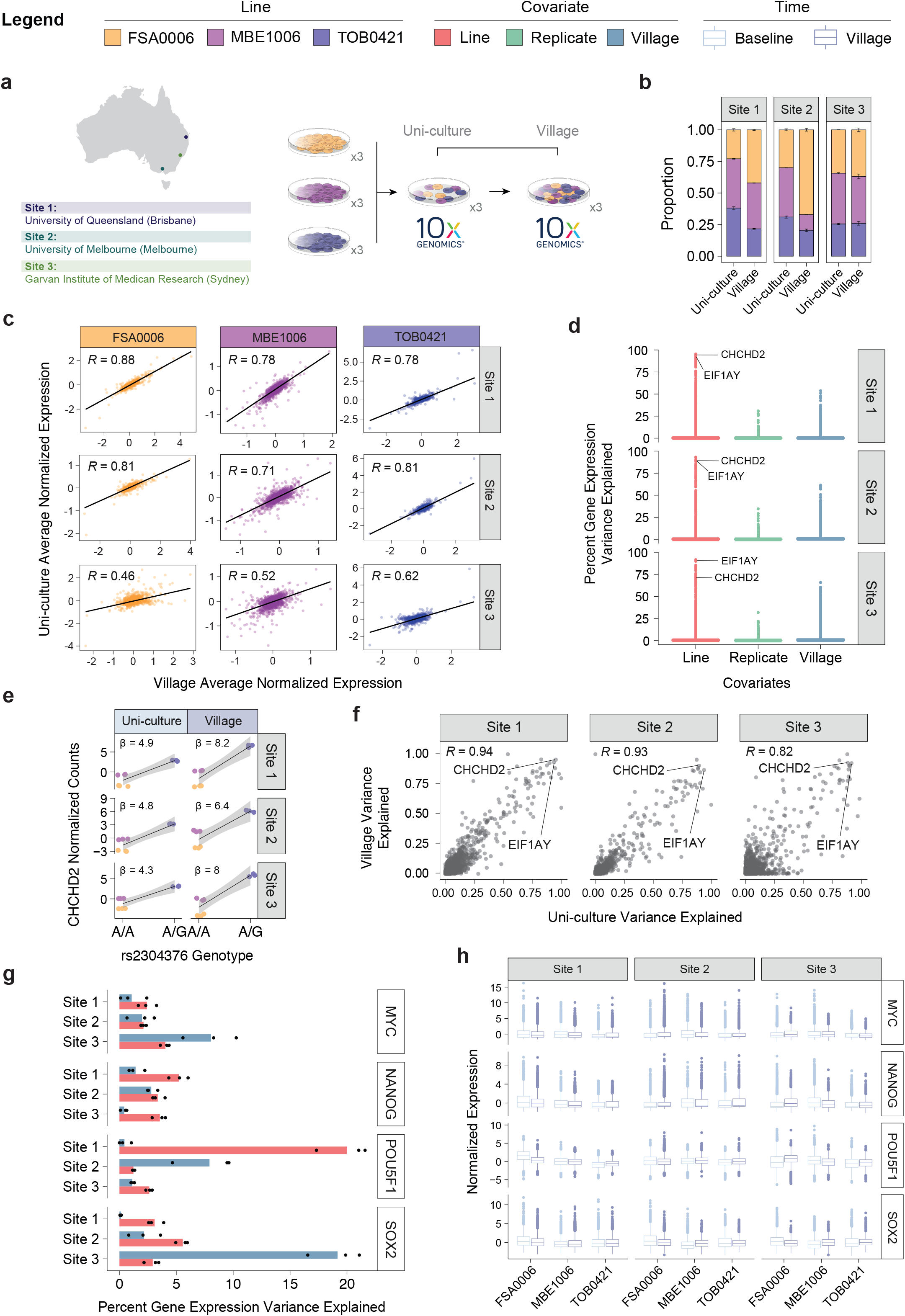
Impact of village culturing system. **a)** Experimental design to test impact of village culturing systems of hiPSC transcriptional profiles. The same experiment was carried out at each of the three different sites in triplicate. **b)** The proportions of each of the three hiPSC lines at each of the three different sites at baseline and after culturing the cells in a village. The proportions between baseline and village are different at Sites 1 and 2 but the same at site 3. **c)** Pearson correlation between the expression profiles of each hiPSC line at each Site at baseline and after village culturing demonstrates a strong relationship between the two conditions (0.46 ≤ R ≤ 0.88). **d)** The variance in gene expression explained by the hiPSC line, replicate and village status demonstrates that the hiPSC line explains a larger percentage of the variance than the replicate or village status. **e)** The previously-reported eQTL for *CHCHD2* demonstrates a strong and consistent effect across different Sites and village status which is even slightly stronger for the cells cultured in a village as compared to when they were cultured separately. **f)** The variation in gene expression that can be explained by the hiPSC lines is highly correlated between baseline and village samples at each site (Pearson correlation: 0.82 ≤ R ≤ 0.94). **g-h)** The variance of important stem cell markers explained by the hPSC lines was relatively small ≤ 5% and was not impacted by the variance explained by the village status. hiPSC: human induced pluripotent stem cell.

### Transcriptional profiles are unaffected by village culturing

To evaluate whether cell signaling companion hiPSC lines in a village alter the transcriptional profiles of each individual line we compared the transcriptome profiles of each hiPSC line between uni- and village cultures. We observed strong correlation of the transcriptional signatures between uni-culture and village samples for each hiPSC line (Pearson 0.88 ≥ R ≥ 0.46, *P* < 2.2×10^−308^; **Figure 2c** and **Table S2**).

To evaluate the effect of culturing hiPSC lines in a village on each gene, we utilized a negative binomial model framework to estimate the percentage of variance of the expression of each gene that was explained by either the line, replicate, or village (**Figure 1b**). Across all three sites, we observed that the hiPSC line explains the largest percentage of the gene expression variance (*i*.*ei*., inter-line variation; **Figure 2d**). This is supported by the observation that genes with the highest variance explained by the hiPSC lines included Y chromosome genes (*e*.*g*., *EIF1AY* - **Table S3**). Similarly, genes with larger inter-line variation included genes that have been reported as expression quantitative trait genes (eGenes) in previous hiPSC eQTL studies (*e*.*g*., *CHCHD2*)^21,22^. These results suggest that culturing hiPSC lines in a village system does not significantly alter the transcriptome of each hiPSC line.

Since the variation in *CHCHD2* expression was largely described by the hiPSC lines and was previously described as an eGene, we next investigated whether we could detect the reported eQTL^22^ with our data. Indeed a previous eQTL between the single nucleotide polymorphism (SNP) rs2304376 was consistently associated with *CHCHD2* expression (4.3 ≤ β ≤ 8.2; **Figure 2e**). As reported previously, the reference A allele was associated with lower expression compared to the alternate G allele. This demonstrates that line-specific effects such as eQTLs can be consistently detected using village culture systems.

In order to more directly test whether the village culture system impacts inter-line gene expression variation, we tested the correlation between the gene expression variance explained by uni-cultured and village samples. The percent of the variance of gene expression explained by the hiPSC line was highly consistent between the baseline and village samples (0.82 ≤ R ≤ 0.94; **Figure 2f**) which demonstrates that cell signaling in village cultures does not significantly alter unique hiPSC line transcriptional profiles.

Finally, we evaluated whether the line and village effects were consistent across sites by estimating the variance of gene expression explained by the hiSPC lines (*i*.*e*., inter-line variation) or explained by the village status between sites. The relationship for the gene expression variance explained by the village (0.41 ≤ R ≤ 0.66; **Figure S1a**) was less than the relationship for the gene expression variance explained by the hiPSC lines (0.88 ≤ R ≤ 0.92; **Figure S1b**) These results indicate that the impact of culturing hiPSC lines in a village system on the cellular transcriptome is largely stochastic - indeed the mean percentage of variance explained by the village effect was only 1.1%. Conversely, the effect of the hiPSC line on gene expression variation is non-random and is consistently detected at independent sites which demonstrates that different laboratories using the same hiPSC lines for village experiments can produce the same results.

### Consistency of hiPSC line effects is not altered in villages

The correlation of inter-line expression correlations were similarly high between uni-cultured and village samples (0.82 ≤ R ≤ 0.94; **Figure 2f**) as well as between samples from different sites (0.88 ≤ R ≤ 0.92; **Figures S1b**). However, it is difficult to detect whether the correlation is consistently high across all percentages of variance explained by the hiPSC lines (*i*.*e*., at 1% vs at 80% variance explained). Therefore, we next interrogated the relationship of genes by assessing the correlation of genes in windows of 100 - starting with the 100 genes with the lowest inter-line variation and sliding the window by one gene until the 100 genes with the highest inter-line variation were correlated (**Figure 1c**).

To compare the results of the between-site and between-village correlation results, we identified the percent variance explained by the hiPSC lines at which a good correlation was achieved (Spearman ⍰ > 0.5). The percent variance explained by the hiPSC lines at this threshold (Spearman ⍰ > 0.5) was not significantly different (P ≥ 0.82) between the inter-village correlations (9.3% - 30.2%) and the inter-site correlations (uni-culture: 12.0% - 23.0%, village: 13.9% - 23.4%; **Figure S1c-e**). These data demonstrate that 1) genes whose variance is only minimally explained by the hiPSC lines are stochastic and 2) the transcriptional differences observed between baseline and village samples are comparable to the differences observed between sites. Therefore, any variation in inter-line gene expression effects observed in village samples is reflective of normal, technical variation.

### hiPSC line effects of pluripotency genes are not impacted by villages

It is important to understand the impact (if any) that the village culture system could have on gene expression denoting pluripotent state. We identified that, on average, a small percentage (≤ ∼5%) of gene expression variance was explained by the hiPSC lines for common stem cell markers such as *MYC, NANOG, SOX2* and *POU5F1*^*5*^ (**Figure 2g** and **Figure S1f**). Even in scenarios where the village status explains a large percentage of the gene expression variation, the gene expression variation attributed to the hiPSC lines is highly consistent with the other sites (*i*.*e*., *SOX2* at Site 3; **Figure 2g-h** and **Figure S1f**). Importantly, these data suggest that even when the village status impacts pluripotent gene expression, the hiPSC line effect is still detectable and consistent.

### Impact of village culture system in cryopreserved samples

Since some experimental designs require cryopreserving cells at different time points, we investigated village effects comparing fresh and cryopreserved uni-culture and village culture samples (**Figure 3a**).

**Figure 3:**
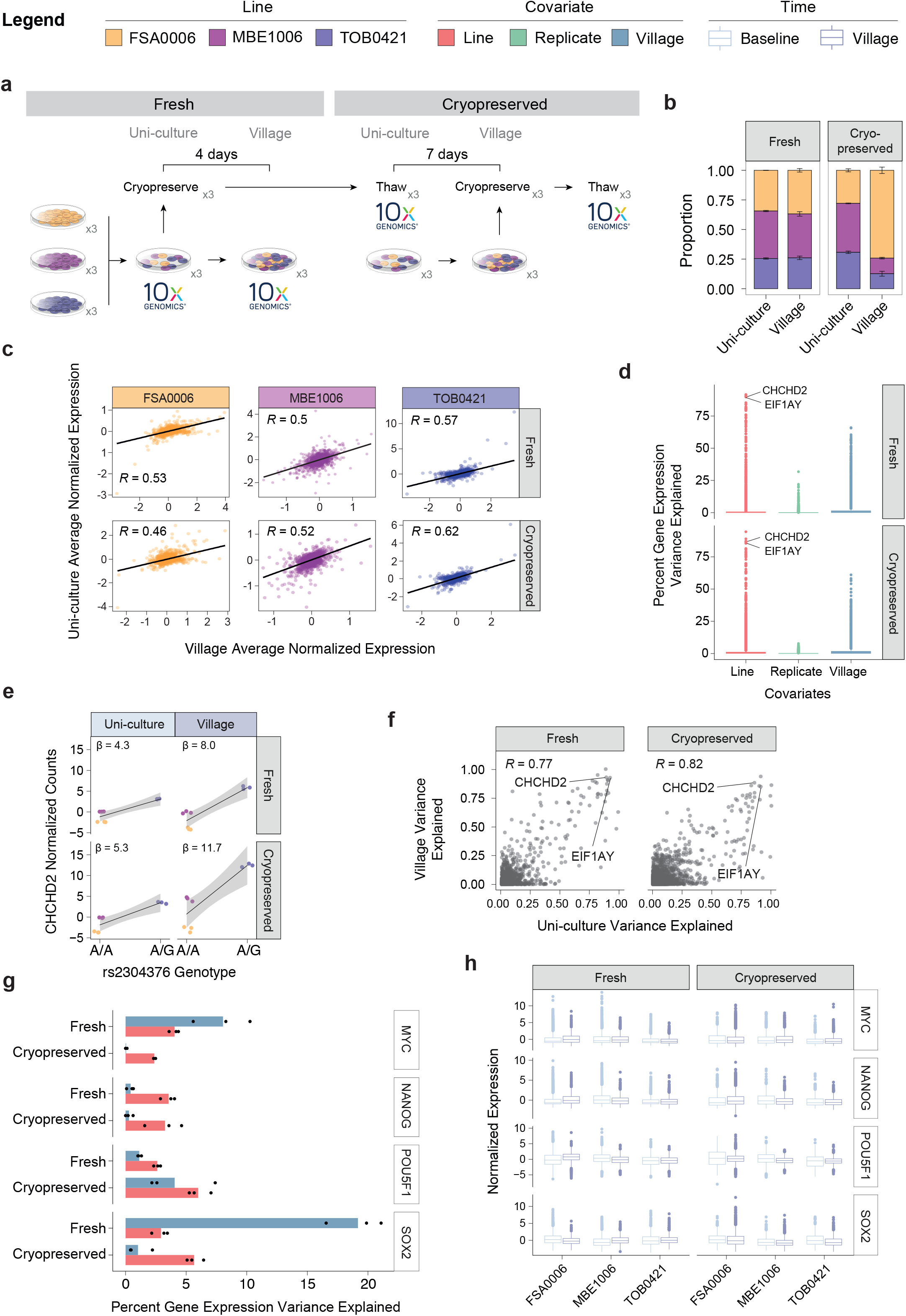
Impact of village culturing system on cryopreserved samples. **a)** The experimental design to test the effect of the village culturing method with cryopreservation - only carried out at Site 3. **b)** The proportion of each hiPSC line was consistent for the fresh but not the cryopreserved samples **c)** The expression profiles of each hiPSC line for fresh and cryopreserved samples are well correlated (Pearson correlation: 0.46 ≤ R ≤ 0.62). **d)** The expression variance explained by the hiPSC lines, the replicate and the village status shows that the hiPSC line explains the largest percentage of the gene expression variance. **e)** Testing for the previously reported eQTL between the rs2304376 SNP and *CHCHD2* expression demonstrates a strong relationship between the SNP genotype and *CHCHD2* expression level. Further, the effects are consistently higher for hiPSC lines cultured as a village than when cultured separately. **f)** The variance observed in gene expression was strongly correlated between baseline and village sample for both fresh and cryopreserved samples (Pearson correlation: 0.77 ≤ R ≤ 0.82). **g-h)** The hiPSC lines explained a small ≤ ∼5% of the variance of important stem cell markers which was unaffected by the village status.

### Proportions of hiPSC lines following village culturing following cryopreservation

Since village culture systems are only effective if cells from all hiPSC lines included in the culture can be detected, we first investigated whether we could detect all three hiPSC lines in our village samples by demultiplexing the samples and identifying the proportion of cells from each hiPSC line. We were able to identify cells from each hiPSC line and found that the proportions of the hiPSC lines when the village was generated for both fresh and cryopreserved samples were similar, indicating that the cryopreservation process does not significantly affect cells from specific lines. In addition, the proportions between the fresh uni-cultured and village-cultured samples were not significantly different (*P*=0.23). However, the proportions of each hiPSC line were different between the cryopreserved uni- and village cultured samples (**Table S4**). The cryopreserved results were consistent with the fresh village sample distributions from sites 1 and 2 (**Figure 2b**) which were plated at similar densities. This reinforces the importance of hiPSC growth rates when choosing compatible lines for hiPSC villages.

### Transcriptional profiles are unaffected by village culturing following cryopreservation

Next, we compared the transcriptional profiles of each of the hiPSC lines that were uni- and village cultured. We found that the expression profiles were well correlated (Pearson R ≥ 0.46; **Figure 3c** and **Table S5**). Further, the inter-line variation explains the largest percentage of gene expression variance (**Figure 3d**). Again, the genes that demonstrated large inter-line variation included Y chromosome genes (*i*.*e*., *EIF1AY*) and genes previously reported as eGenes in hiPSCs (*i*.*e*., *CHCHD2*)^21,22^.

Similar to the results from Phase I, the percentage of gene expression variation that was explained by the hiPSC lines was highly consistent between the uni- and village cultured samples (0.77 ≤ R ≤ 0.82; **Figure 3f**). However, the contribution of the village status to gene expression variance was inconsistent between the fresh and cryopreserved conditions (*R* = 0.49; **Figure S2a**), from which we conclude that the variation explained by the village status was relatively stochastic. Conversely, the variance explained by the inter-line variation was highly consistent between cryopreserved and fresh conditions (*R* = 0.91; **Figure S2b**). The genes whose variance was largely explained by the hiPSC lines included the Y chromosome gene *EIF1AY* and the previously reported eGene *CHCHD2*.

We then tested for the previously reported eQTL between the rs2304376 and the *CHCHD2* gene^21,22^ in these hiPSC lines and observed an effect consistent with that previously reported - the A allele was associated with lower expression of *CHCHD2* compared to the G allele (4.3 ≤ β ≤ 11.7; **Figure 3g**). This effect was highly consistent across cryopreservation and village samples. These results support the conclusion that culturing hiPSC lines in a village model is not likely to impact the ability to identify line-dependent transcriptional effects such as eQTLs.

### Consistency of hiPSC line effects is not altered in villages following cryopreservation

The gene expression variance explained by the hiPSC lines was highly correlated between baseline and village samples (0.77 ≤ *R* ≤ 0.82; **Figure 3f**) as well as between fresh and cryopreserved samples (*R* = 0.91; **Figure S2b**). However, it is not clear if that correlation is consistently high across all percentages of gene expression variation explained by the hiPSC lines (*i*.*e*., whether the correlation has the same strength at 1% and 90% of gene variance explained). In order to address this question, we assessed the same sliding window correlation approach used in Phase I. Each window included 100 genes and was moved by one gene to the next window until the 100 genes with the highest inter-line variance (**Figure 1c**).

Assuming that the pairs of samples between fresh and cryopreserved samples at the same time (*i*.*e*., uni- or village culture samples) are representative of normal variation between samples, we can test whether the inter-culture method variation (*i*.*e*., uni-vs village culture samples) is reflective of normal variation between samples. To test this, we compared the variance explained by the hiPSC lines that achieved a good correlation (Spearman ⍰ > 0.5) using the sliding window approach (**Figure 1c**). Indeed, we observed that the inter-time and inter-cryopreservation samples were not significantly different from one another (Wilcox Test, *P* = 0.21; **Figure S2c-e**). These data indicate that variation between baseline and village samples are reflective of normal sample-to-sample variation and that genes that have small inter-line variation are relatively stochastic.

### hiPSC line effects of pluripotency genes are not impacted by villages culture following cryopreservation

We next interrogated the expression variance explained for pluripotency genes (*POU5F1, SOX2, NANOG* and *MYC*, among others). Consistent with Phase I results (**Figure 2h-i** and **Figure S1b**), we observed that only a small percentage of the total pluripotency gene variances could be explained by the lines and that this effect was consistent between fresh and cryopreserved samples. This was true even when the impact of the village status on gene expression variation was large - for example the fresh *SOX2* variance (**Figure 3h-i** and **Figure S2h**).

Taken together, these results suggest that cryopreservation of villages does not significantly alter the proportions of each hiPSC line or their unique transcriptional profiles and suggests that village systems are indeed appropriate for population genomic studies.

### Dynamic effects of hiPSC line across cell state pseudotime

From our results we conclude that culturing cells in village conditions does not significantly alter inter-line variation in gene expression across cells. However, our results until this point do not take into account the potential variation in cell states. We next sought to determine the variance in gene expression explained in different cell states - for example differences in pluripotent potential or cells spontaneously differentiating.

Using RNA velocity, we first positioned cells based on their estimated pseudotime trajectory (**Figure 1d**). As expected, we observed that the pseudotime landscape was strongly defined by pluripotency of the cells. The cells that had lower pseudotime values were more pluripotent and the cells with higher pseudotime had markers for spontaneous differentiation into the neural and epidermal ectoderm lineages (**Figure 4a-e**). To interrogate contributions to gene expression variation across the pseudotime, we split the pseudotime into quintiles (**Figure 1d** and **Figure 4f**) and assessed the variance explained by the hiPSC line, village status and replicates.

**Figure 4:**
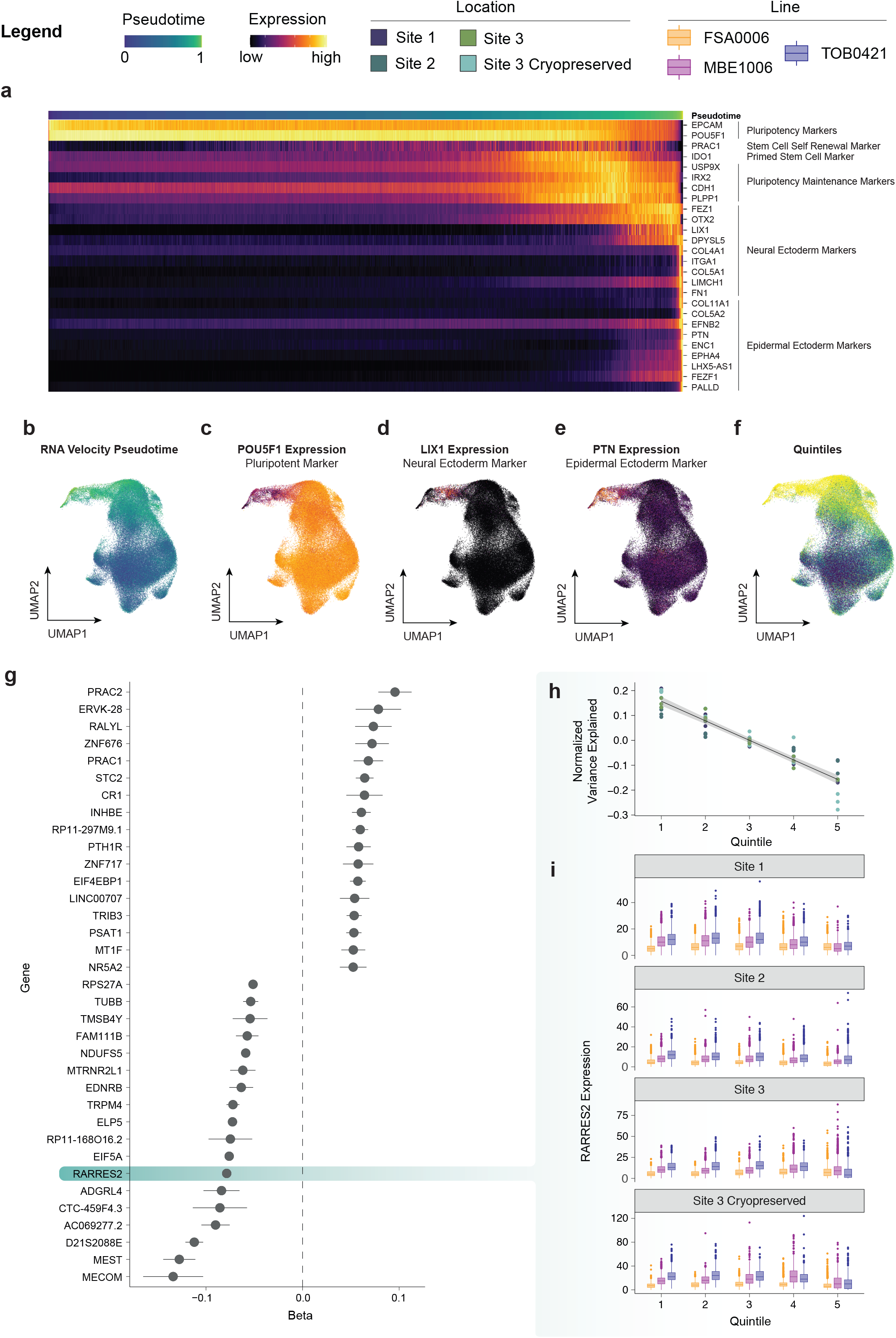
Dynamic Variance Explained Across Stem Cell Pseudotime. **a)** Pseudotime was defined by pluripotency markers and ectodermal markers with the lower pseudotime corresponding to spontaneously differentiated cells. **b)** The pseudotime projected onto the UMAP of all cells. **c-e)** Markers representative of the three groups defining the pseudotime: pluripotency (*POU5F1*), Neural Ectoderm (*LIX1*) and Epidermal Ectoderm (*PTN*). **f)** The quintiles used for dynamic variance explained detection projected onto the UMAP. **g)** The top dynamic variance explained genes. **h-i)** For example, *RARRES2* demonstrates a consistent decrease in variance explained across the different quintiles described by each Site.

### Gene expression variance explained by hiPSC lines is unaffected by villages across pseudotime

Similar to our previous findings, the line effect explains a larger percentage of variance than the village or replicates (**Figures S3a**) and the relationship of variance explained between different sites for each quintile was stronger for the line than the village status (*P* < 0.001; **Figure S3b**). Further, we show that the correlation of the variance explained between quintiles was larger for neighboring quintiles (ie Q1 and Q2) than for distant quintiles (ie Q1 and Q5; **Figure S3c-d**).

### hiPSC line effect is dynamic across pseudotime

The stronger similarity of inter-line variance between neighboring quintiles (**Figure S3c**) suggests that the gene expression variance explained by the hiPSC lines is dynamic, although consistently changing across pseudotime. To test this hypothesis, we used linear models to estimate the relationship in the gene expression variance explained across quintiles - correcting for the different conditions (site 1, site 2, site3 and site 3 Cryopreserved; **Figure 1d**). We identified 1,965 significant dynamic line effects - inter-line variation that consistently decreases or increases across pseudotime (FDR < 0.05, **Table S6**). Thirty-five of those significant dynamic effects had an effect size (β) between -0.05 and 0.05 (**Figure 4g**). Importantly, none of those significant dynamic genes were pluripotency genes - indicating that any inter-line variation of pluripotency genes is consistent across pseudotime.

The 35 genes with large dynamic effect sizes included the *RARRES2* gene - a retinoic acid receptor responder that is an important chemoattractant for immune response^23^. The interline variation was larger for the more pluripotent quintiles (Q1) than the spontaneously differentiated quintiles (Q5; **Figure 4h**). This effect was highly consistent across different sites and replicates (**Figure 4h-i**), which confirms that village culture systems do not significantly alter transcriptional profiles - even across pseudotime or different cell states.

*RARRES2* has previously been described as an eGene in hiPSCs^21^ and a dynamic eGene dependent on aryl hydrocarbon receptor (AHR) activation^24^. The association between the dynamic eQTL rs2108852 SNP genotype and *RARRES2* expression in the pluripotent quintile (Q1) of the hiPSCs was consistent with the previously reported effect where the alternative C allele was associated with higher expression of *RARRES2*. Further, this effect (β) continuously decreased with each sequential quintile (**Figure S5e**). rs2108851 is also in high linkage disequilibrium (R^2^ = 0.86) with the eQTL SNP previously described in hiPSCs (rs11766288), and demonstrates the same allelic direction of effect^21^. These results support the conclusions that sensitive, dynamic differences in the inter-line expression variation in hiPSC lines can be effectively detected using village culturing systems.

Collectively, these data demonstrate that village culture methods can be effectively used for population genomic studies with hiPSC lines. We have also demonstrated that cryopreservation does not alter the transcriptional profiles or hiPSC line proportions and even sensitive dynamic effects can be reproduced with village systems.

## Discussion

Advances in human genetics, stem cell biology, and single cell technologies have given rise to the convergence of these research domains that allows better evaluation of the complexity of human genetic regulation. Population genomic studies using hiPSCs and hiPSC-derived cells have steadily increased in sample size and frequency as more hiPSC lines have become available and methods for culturing hiPSCs and hiPSC-derived cells have developed. However, maintaining hiPSCs and hiPSC-derived cells is still expensive and time consuming which compromises the ability to apply large-scale population genomic studies with consistency.

To date, the majority of population genetics hiPSC studies have been conducted using bulk sequencing methods - which means that the transcriptomes of all the cells in a sample are combined and assayed together as one. These studies have provided significant new knowledge on the role of genetic variation in genome regulation, although by using bulk sequencing approaches, they effectively remove any heterogeneity in a sample which is important for cell type-specific and context-specific effects. We argue that in contrast, single cell technologies provide a powerful solution for this challenge, as individual cells are assayed separately and context-specific effects can be interrogated. Indeed, a few recent studies have demonstrated that single cell and deconvolution methods applied to hiPSCs and hiPSC-derived cells can be used for detection of pQTLs^11^ and eQTLs, and identify context-specific effects^2–4,8^.

Studies that have used village culture systems coupled with scRNA-seq have employed computational demultiplexing approaches to effectively obtain RNA measures for each cell from each hiPSC line^2,8^. Those studies have identified context-dependent eQTLs, but have not assessed the potential impact of cell signaling in the village culture system, and whether it alters the transcriptional profiles of companion lines. They detected a lower number of eQTLs per cell type than anticipated for the sample sizes which could have been partially due to introduction of non-genetic variance as a result of the village model^2,8^. Therefore, to confidently use village models for future population-scale experiments, there was a critical need to thoroughly assess whether using a village culture system would alter the transcriptional profiles of the cells from each hiPSC line. Our experimental procedure was specifically designed to address that question and provide a roadmap to village culture experimental design.

Our results demonstrate that, while it is important to consider hiPSC line growth rates, village experimental culture systems do not alter the unique transcriptional profiles of each hiSPC line and are therefore an applicable system for population-scale experiments. Importantly, we were able to detect each hiPSC line in all samples, and consistently identify line-dependent variation. Transcriptional profiles were highly consistent in villages and at different sites which also demonstrates that village experiments could, theoretically, be conducted across multiple sites. In addition, cryopreservation had no detectable effect on transcriptional profiles regardless of whether they were cryopreserved at the time of forming the village or after being cultured in a village. Therefore, we conclude that line-dependent studies like QTL studies can effectively be carried out by using village culturing designs.

The advantages gained from using single cell data to obtain purer cell subtypes and from leveraging village culture systems to increase throughput are extensive. In our opinion, these advantages significantly outweigh the possibility of not detecting a small effect size QTL for studies in hiPSCs. Village systems - paired with single cell technologies - promise to revolutionize the field of population genomics of gene regulation.

## Materials and Methods

### Ethics

The research carried out in this study was in accordance with the Declaration of Helsinki and approved by the Human Research Ethics committees of the University of Melbourne (1545394), the Garvan Institute of Medical Research (ETH01307) and the University of Queensland (2015001434).

### hiPSC Cell Culture

1mL aliquots of each of the hiPSC lines FSA0006, MBE1006 and TOB0421^20^ (passage 18; **Supplementary Table S3**) were thawed at each site on the same day and plated in StemFlex media (Life Technologies; Catalog Number: A3349401; Lot Number: 2093181) complete with StemFlex Supplement (Life Technologies, Catalog Number: A33492-01; Lot Number: 2090179) with Rock inhibitor Y-27632 (10µM final concentration; Stem Cell Technologies; Catalog Number: 72304) on Costar non-treated 6 well polystyrene plates (Catalog Number: 3736; Lot Number: 30417038) coated with Vitronectin XF (Stem Cell Technologies; Catalog Number: 07180; Lot Number: 18B87584) diluted in CellAdhere Buffer (Stem Cell Technologies, Catalog Number: 07183; Lot Number: 18M979058). Cells were subcultured once a week with ReLeSR for two weeks (Stem Cell Technologies; Catalog Number: 05872) before equal numbers of the hiPSC lines were combined and plated together. The same lot number of all reagents were used between the three locations (University of Queensland in Brisbane, the University of Melbourne in Melbourne and the Garvan Institute of Medical Research in Sydney).

### scRNA-seq Capture

Totalseq-A antibodies (Biolegend) were used to hashtag hiPSC pools from each different location (University of Queensland, Garvan Institute of Medical Research and University of Melbourne) before combining and superloading onto the 10X Genomics Chromium Controller (10x Genomics) to capture single cells. Briefly, 1×10^6^ cells from each replicate at each site were centrifuged at 300g for three minutes and the supernatant was discarded. The cell pellets were resuspended in 100µL cold Fluorescence-Activated Cell Sorting (FACS) buffer (phosphate buffered saline [PBS] with two percent fetal bovine serum [FBS]). Then, 2µL of Totalseq-A hashing antibody was added and the cells were gently pipetted to mix before incubating for 20 minutes on ice. Cells were then washed twice with FACS buffer by centrifuging at 300g for 5 minutes, discarding the supernatant and resuspending the cell pellet in 100µL cold FACS buffer. Cells were briefly stained with 4′,6-diamidino-2-phenylindole (DAPI) before using flow cytometry (BD FACS Aria, 100um nozzle, 4-way purity mode, temperature controlled) to sort and capture live single cells. Cell pools contained one sample from each site. Trypan blue was then used to assess the pool viability (> 75% viable). Approximately 32,000 cells were loaded onto the Chromium Single Cell Chip B (10X Genomics; Catalog Number: PN-1000073) in order to capture 20,000 single cells with the Gel Bead Kit V3.0 (10x Genomics; Catalog Number: PN-1000076).

GEM generation, barcoding, cDNA amplification, and library construction were performed according to the 10x Genomics Chromium User Guide (CG000183). Libraries were prepared with the Single Cell 3’ V3.0 Library and Gel Bead Kit (10x Genomics; Catalog Number: PN-1000077 and PN-1000078).

### scRNA-seq Read Alignment

The 10x Genomics Cell Ranger Single Cell Software Suite (version 3.1.0) was used to process the 3’ single cell RNA-seq libraries (chemistry v3). Raw base calls were used to demultiplex the multiplexed pools, which were then mapped to the GRCh38-1.2.0 genome (Ensembl release 84) using STAR for each pool independently.

### scRNA-seq Demultiplexing and Doublet Detection

The lines in the scRNA-seq pools were demultiplexed using SNP genotype demultiplexing with *Popscle Demuxlet v0*.*1-beta*^*13*^, *Popscle Freemuxlet v0*.*1-beta*^*25*^, *scSplit v1*.*0*.*1*^*14*^, *Souporcell v1*.*0*^*15*^, and *Vireo v0*.*4*.*2*^*12*^. These methods both demultiplex the different hiPSC lines in the pools and identify doublets between two different lines. The recommended guidelines were followed for each of the demultiplexing softwares as briefly described.

### Popscle Demuxlet

*Popscle pileup* was used to identify the single nucleotide variants (SNVs) in the pool. Then, *Demuxlet* was run with reference genotypes for each hiPSC line in the pool using a genotype error coefficient of 1 and genotype error offset rate of 0.05 and default options for all other parameters.

### Popscle Freemuxlet

*Popscle pileup* was used to identify the single nucleotide variants (SNVs) in the pool followed by *Freeuxlet* executed with default parameters.

### scSplit

Low quality and duplicated reads were removed before using freebayes to classify high quality SNVs in the dataset. The resulting bam and vcf were used for *scSplit* using default options and the -n 3 option to provide the number of hiPSC lines in the pool.

### Souporcell

*Souporcell* was run using the *souporcell_pipeline*.*py* script with known variant locations from the reference imputed SNP genotypes that overlapped gene exons using the *--common_variants* parameter and all other default parameter options.

### Vireo

Model 1 of cellSNP v0.3.2 was used to identify allele frequencies at the locations of the common variants (MAF = 0.1) in the genotyped reference genotype file for the three hiPSC lines. The resulting pileup was filtered for SNP locations that were covered by at least 20 UMIs and had at least 10% minor allele frequency across all droplets. Vireo version 0.4.2 was then used to demultiplex the droplets in the pool using reference SNP genotypes and indicating the number of individuals in the pools.

### Scrublet

*Scrublet* was used to identify transcription-based doublets that included two cells from different cell types in. Scrublet was implemented in *python v3*.*6*.*3* per developer recommendations with at least three counts per droplet, three cells expressing a given gene, 30 PCs and a doublet rate based on the following equation:

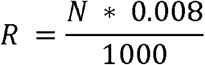

where *N* is the number of droplets captured and *R* is the expected doublet rate. Scrublet was assessed at four different minimum numbers of variable gene percentiles: 80, 85, 90 and 95. Then, the best variable gene percentile was selected based on the distribution of the simulated doublet scores and the location of the doublet threshold selection. In the case that the selected threshold does not fall between a bimodal distribution, those pools were run again with a manual threshold set.

### Hashtag Demultiplexing

Hashtag demultiplexing^17^ was used to identify cells from each location and doublets that included cells from two different locations.

Droplets that were classified as singlets by at least four of the SNP-based demultiplexing or transcription-based doublet detecting softwares as well as the hashtag demultiplexing and were classified as the same hiPSC line by at least three of the SNP-based demultiplexing softwares were retained for downstream analysis. All other droplets were excluded.

### Quality Control

Droplets were considered outliers and excluded from further analysis if they were more than four median average deviations (MAD) from the mitochondrial percent median or contained less than 1,750 total genes. This resulted in 88,927 high quality single cells to be used for downstream analysis. Cyclone from the scran package v1.4.5^26^ was used to detect the cell cycle state of each cell. The quality control metrics for these high-quality single cells for the fresh samples (**Figure 2a**) are presented in **Figure S4** and the cryopreserved experiment samples (**Figure 3a**) are presented in **Figure S5**.

Data were normalized with a regularized negative binomial regression and variance stabilized with Pearson residuals as previously described^27^. Expression data were also corrected by cell cycle status, mitochondrial percent and ribosomal percent.

### Correlation of Transcriptional Profiles

The mean expression of each gene for each hiPSC line at each site was compared between baseline and village samples with a Pearson correlation.

### Covariate Contribution to Gene Variance

The proportion of the variance explained by the hiPSC line, the replicate and the village status for each gene was determined by fitting a negative binomial model for normalized UMI counts of each gene following a method previously described^28^. Briefly, normalized UMI counts were fit as the dependent variable of a negative binomial model with the independent variables fit as random effects. The intercept and variance were then used to estimate the within-cluster variance 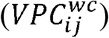 for each independent variable (equation 1) where 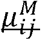 is the marginal expectation defined in equation 2.

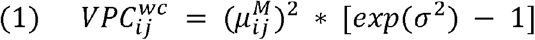

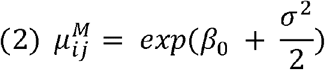

### Sliding Window Methods for Identify Consistency Gene Variance Explained

To identify the percent variance explained by hiPSC line effects that were consistent between conditions (between different sites or different times) we used a sliding window approach. Each window included 100 genes and was shifted by one gene starting with the 100 genes with the lowest variance explained by the hiPSC lines and ending with the 100 genes with the highest variance explained by the hiPSC lines. The spearman correlation coefficient was calculated for each pair of sites at each time (baseline or village) and for each pair of times at each same site.

### RNA Velocity Pseudotime

Pseudotime was estimated using RNA velocity implemented with the *scvelo*^*29*^ package (v0.2.3) to estimate latent time of all single cells. First, sequence reads overlapping spliced and unspliced read count matrices were prepared using *velocyto*^*30*^ (v0.17.17). Cells that had less than 1000 unspliced counts and genes that were expressed in less than 20 cells and had less than 10 unspliced counts were filtered and removed. The batch effects were normalized using *pycombat*^*31*^ (*Combat v0*.*3*.*0*) using a proposed approach^32^. Briefly, the spliced (*S*) and unspliced (*U*) counts were combined to create the total counts matrix (*M*; equation 3). An additional matrix (*R*) was also constructed to aid in deriving the corrected spliced (*S*_*b*_) and unspliced (*U*_*b*_) matrices following batch correction of *M*(equation 4). Then M was corrected for site, hiPSC line and village status batch effects using *pycombat*. Following batch correction, batch corrected splicted (*S*_*b*_) and unspliced (*U*_*b*_) matrices were derived using the Rmatrix (equations 5& 6).

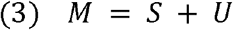

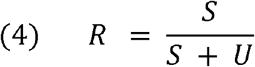

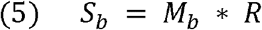

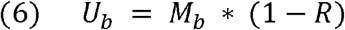

Those matrices were then used to calculate the dynamical RNA velocity latent time with the *scvelo* package.

### Integrating Conditions for Visualization

Cells from each cell line in each village status and at each site were integrated for visualization using the reciprocal principal component analysis (RPCA) method implemented with the *Seurat* package (v4.0.0)^33^.

## Acknowledgments

We are grateful to the participants who provided samples to generate the human induced pluripotent stem cells used in this study. We also want to thank Antonio García for the support on figures.

## Data Accessibility

Data are in the process of being made publicly accessible.

