## Supplementary Figures for "Village in a dish: a model system for population-scale hiPSC studies"

**Supplementary Tables**

**Supplementary Table S1: hiPSC line proportion differences.** The proportions of each hiPSC line at each site were tested for significant differences between baseline and village samples with a paired *t*-test.

**Supplementary Table S2: Expression correlation gene expression between baseline and village culture conditions.** The Pearson correlation results for each hiPSC line at each Site.

**Supplementary Table S3: hiPSC Line Metrics.** Basic metrics for hiPSC lines.

**Supplementary Table S4: hPSC line proportion differences for cryopreserved.** The proportions of hiPSC lines at Site 3 for fresh and cryopreserved samples were tested for differences between baseline and village samples with a paired *t*-test.

**Supplementary Table S5: Expression correlation of transcriptional profiles between baseline and village culture conditions for fresh and cryopreserved samples.** The Pearson correlation summary statistics for the correlation between the baseline and village transcriptional profiles for each hiPSC line at each Site.

**Supplementary Table S6: Dynamic gene variance line effects.** The results from the genes whose variance was significantly associated with pseudotime quintiles demonstrating a dynamic effect across pseudotime.

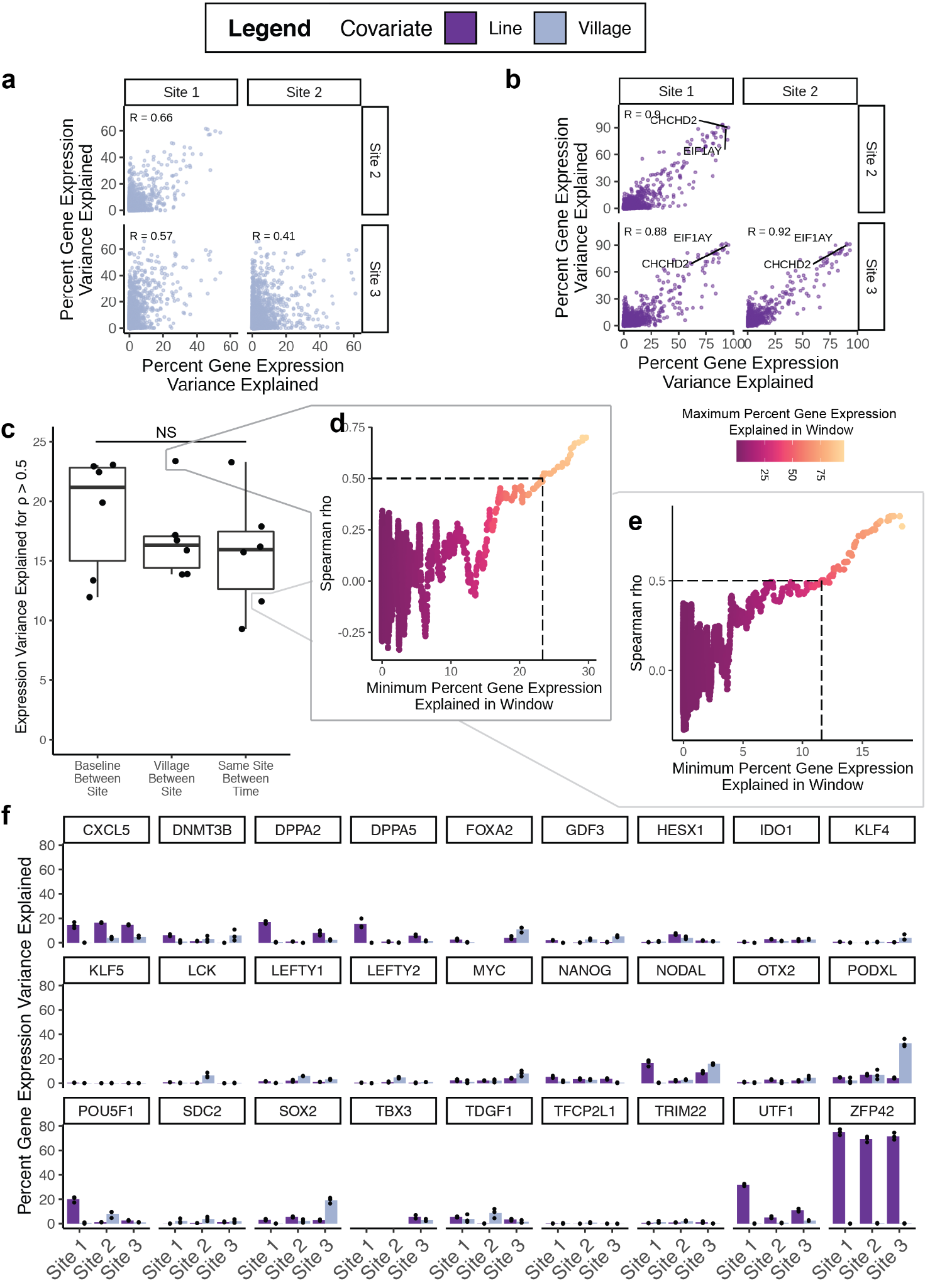

**Supplementary Figure S1: Impact of village culturing systems. a)** The percentage of gene expression variance explained by the village status is not consistent between the sites (Pearson correlation: 0.41 ≤ R ≤ 0.66). **b)** The variance explained by the hiPSC lines was highly consistent between sites which indicates that this effect is non-random and detectable across multiple different conditions (Pearson correlation: 0.88 ≤ R ≤ 0.92). **c)** The expression variance explained by the hiPSC lines when good correlation of the two samples was achieved (Spearman ⍴ > 0.5). There was no significant difference between the different groups (Wilcox Test *P* > 0.05). **d**,**e**) Example plots of the 100 gene window correlation; each point represents a window of 100 genes **f)** The hiPSC lines did not account for a large percentage of the variance explained for most pluripotency genes and was unaffected by the variance explained by the village status.

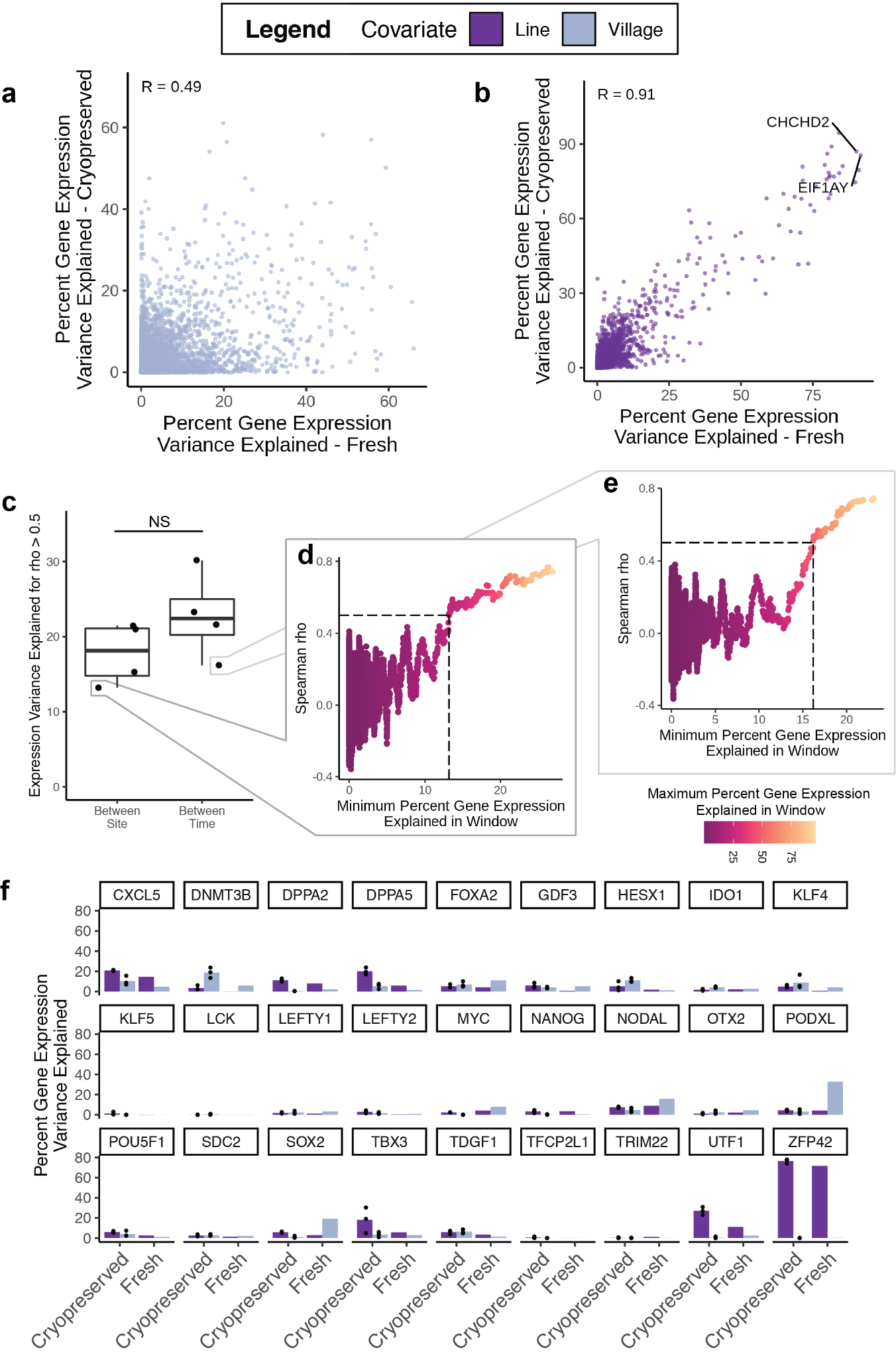

**Supplementary Figure S2: Impact of village culturing system on cryopreserved samples. a)** The percent variance explained by the village status is poorly correlated (R ≤ 0.49) between different sites which indicates that the effect is relatively random. **b)** The percent of the variance explained by the hiPSC line is highly consistent between sites (R = 0.91) which indicates that it is a consistent effect. **c)** The expression variation explained by the hiPSC line at which a good correlation was achieved (Spearman ⍴ > 0.5) was not significantly different (Wilcox Test) for sample pairs at the same site (baseline vs village) compared to the sample pairs between different sites (at the same time). Examples of those paired correlations (**d** and **e**) demonstrate a large amount of noise when only a small percent of the variation in gene expression is explained by the hiPSC line. **f)** The variance of pluripotency genes explained by the hPSC lines (purple) and village status (blue).

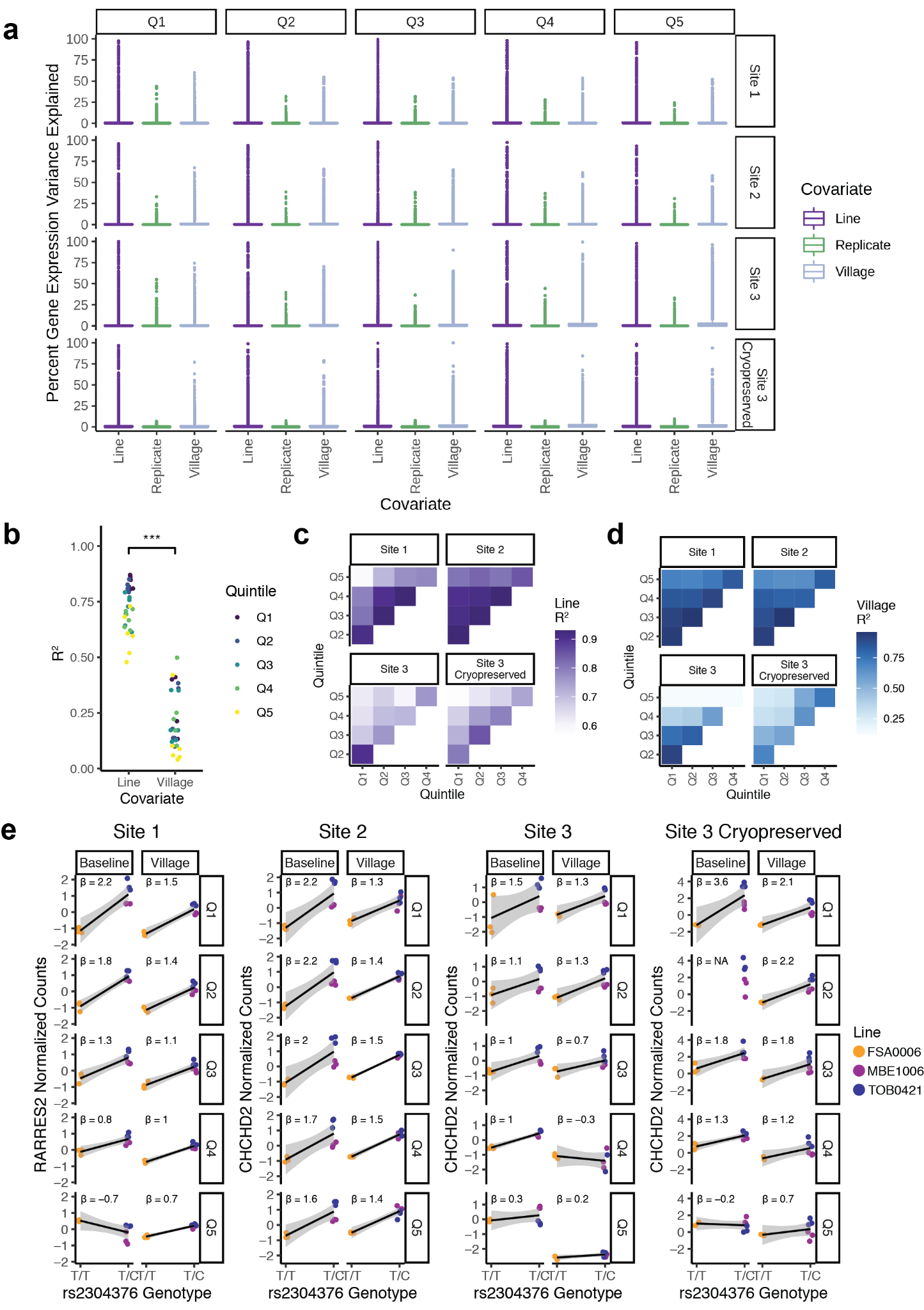

**Supplementary Figure S3: Dynamic variance explained across pseudotime. a)** The variance explained by the hiPSC lines, the replicates and the village status in each of the quintiles at each site. **b)** The correlation of the variance explained by the hiPSC lines in each quintile were significantly larger than the correlation between the variance explained by the village status. **c-d)** The correlation between the variance explained by the hPiSC lines (**c**) and village (**d**) between the quintiles at each site. The quintiles closer to one another demonstrate a stronger relationship than those further from one another. **e)** The dynamic relationship between the rs2304376 SNP and *RARRES2* expression was observed with the sequential decrease in effect size (β) from Quintile 1 to Quintile 5 Sites 1-3 and Site 3 cryopreserved. ***: *P* < 0.001

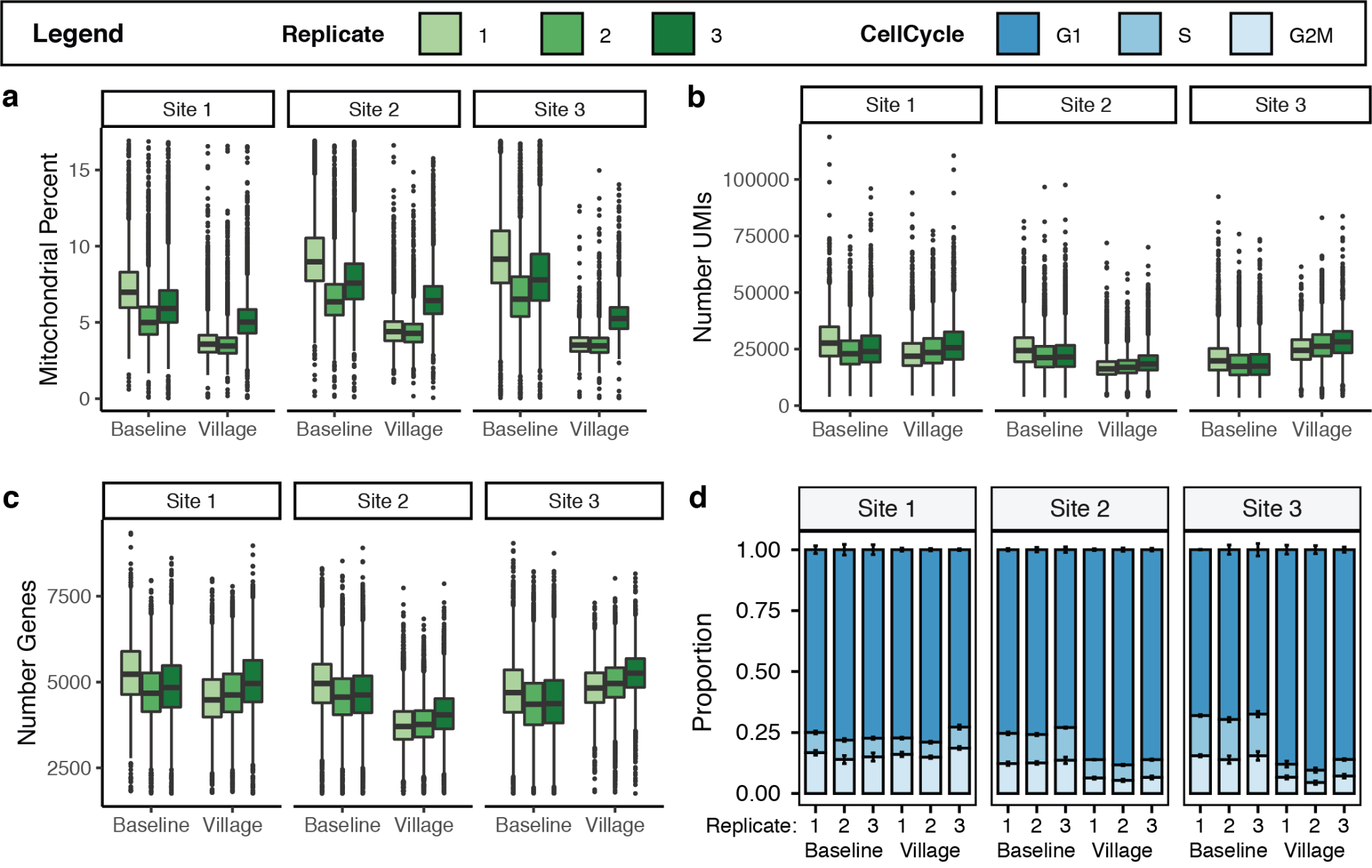

**Supplementary Figure S4: Quality Control Metrics at Three Sites. a)** The mitochondrial percent was different between baseline and village cultured cells but relatively consistent between the sites. **b-c)** The number of UMIs and number of genes detected is relatively consistent between Sites and village status. The proportions of cells in each cell cycle group was relatively consistent between baseline and village culturing conditions for Site 1 but there were more cells in G1 and less in S and G2M phases for Sites 2 and 3.

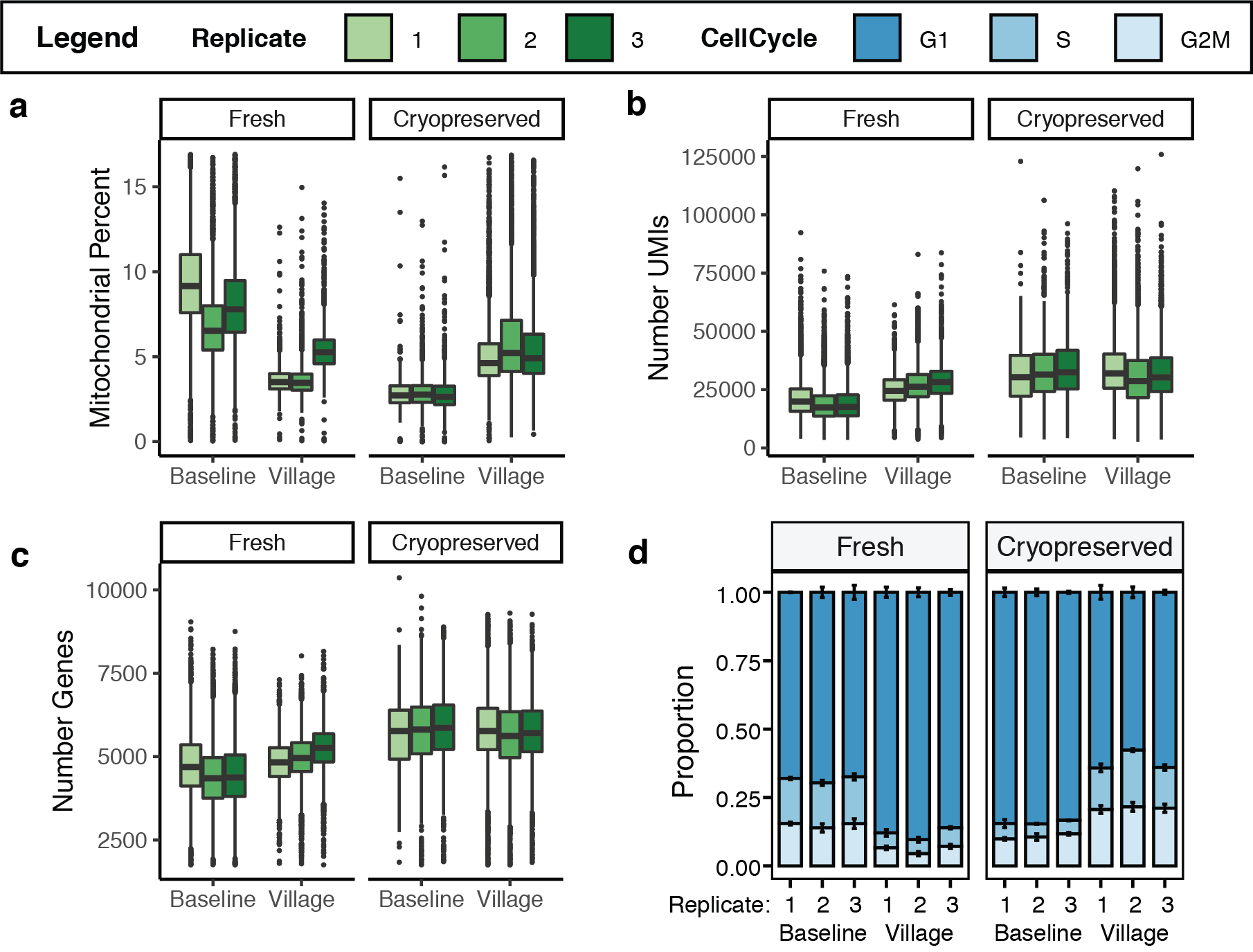

**Supplementary Figure S5: Quality Control Metrics for Fresh and Cryopreserved Samples. a)** The mitochondrial percent were not consistent for cells collected at baseline or village statuses or between fresh and cryopreserved samples. **b-c)** The number of UMIs and genes were relatively consistent between baseline and village samples but were higher in the cryopreserved samples. **c)** The proportions of cell cycle groups were consistent between replicates in each condition but inconsistent between conditions.
